# An empirical attack tolerance test alters the structure and species richness of plant-pollinator networks

**DOI:** 10.1101/2020.01.28.923177

**Authors:** Paolo Biella, Asma Akter, Jeff Ollerton, Anders Nielsen, Jan Klečka

## Abstract

Ecological network theory hypothesizes a link between structure and stability, but this has mainly been investigated *in-silico*. In an experimental manipulation, we sequentially removed four generalist plants from real plant-pollinator networks and explored the effects on, and drivers of, species and interaction extinctions, network structure and interaction rewiring. Our results indicate that cumulative species and interaction extinctions increased faster with generalist plant loss than what was expected by co-extinction models, which predicted the survival or extinction of many species incorrectly. In addition, network nestedness decreased, modularity increased, and opportunistic random interactions and structural unpredictability emerged, which are all indicators of network instability and fragility. Conversely, interaction reorganization (rewiring) was high, asymmetries between network levels emerged as plants increased their centrality. From the experimental manipulations of real networks, our study shows how plant-pollinator network structure has low stability and changes towards a more fragile state when generalist plants are lost.

## Introduction

Interactions are organized in complex networks, and the way these structures react to disturbance is crucial for understanding network functioning, their ability to buffer negative impacts and also for their conservation ^1–4^. This is usually verified with “attack tolerance tests” that assess the functionality of a system after knocking out its important components^5^. In ecology, such tests usually consist in removing all species in one trophic level and then in assessing how many species in another level lost all interactions ^6,7^. So far, in pollination networks, this has been addressed mainly theoretically with numerical simulations that show a higher rate of pollinator extinction when highly linked plants are removed^7–10^. However, these theoretical predictions were not compared to empirical data from similar manipulations, which is urgently needed to assess their reliability^11^.

Manipulative experiments of plant-pollinator networks can illuminate the factors maintaining network stability and the processes of network re-organization. For instance, previous experiments removing only one generalist plant^12,13^ showed that networks are quite stable to this loss, and that other species occupy the role in the network of the removed species. Conversely, when multiple invasive plants are removed, network interaction diversity and generalisation are impacted ^14^, indicating that losing multiple species can strongly affect real networks. Moreover, after disturbance, network stability could depend on the amount of interaction rewiring^18^, i.e. foragers’ ability to use alternative resources after depletion or disappearance of those previously used ^15–17^. Rewiring and the establishment of interactions between plants and pollinators may be regulated by several ecological drivers, such as species trait matching ^19,20^, flower’s rewards ^21,22^ or species abundances ^23,24^. Similarly, it was shown that, in a context of network disturbance, the redistribution of pollinators is constrained by plant traits^25^. Nevertheless, opportunism can prevail over strict interaction rules if foragers, to avoid competition, exploit less rewarding resources^26^. Still, it is unknown how the above-mentioned or similar ecological drivers would rule a perturbed plant-pollinator network.

In this study, we conducted a field experiment in which we sequentially removed several generalist plant species from real networks and investigated the impact on pollinators, their interactions and network structure. We present two alternative expectations that link pollinator foraging strategy and network structure. After plant removal, if foragers will predominantly increase their use of alternative resources (i.e., high rewiring), then network compartmentalization (modularity) will likely decrease, because new interactions might happen with different kinds of resources (i.e., across different compartments)^27^. The other expectation is based on the central position that generalist plants cover in the networks (i.e. hubs^28^). The loss of central nodes, that maintain network cohesiveness and links different modules^3^, would break a network down to isolated subnetworks or compartments following generalist plant removal.

Here, we investigated (a) if the rate of real species and interactions extinction is similar to simulations from established co-extinction models; (b) alterations in the structure of plant-pollinator networks and the rate of interaction rewiring emerging during the plant removal; and (c) what ecological factors mediates these changes.

## Methods

The study included three treatment sites and one control site, located at a mean distance of 2.01±0.95 km from each other, near Český Krumlov, in the Czech Republic (treatments: Site 1 ca 1500 m^2^ in size, 48°49’26.8’’N-14°16’26.2’’E; Site 2, ca 1800 m^2^, 48°49’51.63’’N-14°17’34.12’’E; Site 3, ca 1600 m^2^, 48°49’35.07’’N-14°18’8.2’’E; untreated control: 48°49’26.8’’N-14°16’26.2’’E). Each site was a small grassland with a barrier of trees to likely limit pollinator movements to the surrounding landscape (see details in ^25^). Due to the high mobility of pollinators, we deemed that an experimental design based on small within-site treatment plots would not be appropriate (e.g. ^29^). Hence, the experiment consisted of sequentially removing all inflorescences of the most generalist plant species from the entire surface of the treatment sites, one species at a time until four species were removed (see details in ^25^). We sampled flower-visiting insects in six 10m x 1m transects per site during two days for each experimental phase (before and after each species was removed), and synchronously in the control site. After each “before” phase, flower-visitors were counted and this was used as a proxy of generalization to determine which plant species should be removed next; as detailed in ^25^, this proxy was reliable and in fact we later verified that these plants were visited by the most diverse set of pollinators, similarly to ^13^. We identified all insects to species where possible, otherwise morpho-species were used when necessary (after pre-sorting into families and genera). In addition, we counted the number of flowers or inflorescences of all plant species within transects over the sampling period.

### Species co-extinctions

We compared the rate of pollinator and of interaction extinctions from the field after the removal of each generalist plant to what was expected from two co-extinction models, the Topological (TCM ^7^) and the Stochastic co-extinction models (SCM ^10^). TCM is based on the topology of a qualitative binary network and secondary extinctions (i.e. pollinators) are considered as when a species has no surviving partners after a primary extinction (i.e. plant extinction) has occurred. The SCM uses quantitative data, as it calculates an extinction probability based on the interaction strength between species and the dependency on the interaction (*R*), and allows cascading extinction chains ^10^. Separately for each plant removal stage of the treatment sites, these co-extinction models were triggered by removing the same generalist plant species as the field manipulations, and the number of extinctions were counted. In the SCMs, we ran 10^3^ SCM simulations, and, following^30^, we assigned random *R* values to plants and pollinators.

We counted as extinctions the number of pollinators or the amount of interactions recorded before a plant removal that did not occur after a plant removal, for both the observed networks and the model predictions. To avoid overestimations, in the observed networks we considered (i) as extinct species, those pollinators that had interacted with the plant targeted by removal that were not recorded after the removal, and (ii) as extinct interactions, the difference in the amount of interactions after excluding the species unique to the after phases. In addition, all singletons (i.e. species with interaction abundance of 1) were removed from the observed networks and also from the simulations, to avoid overestimations due to species with extremely small populations and sampling stochasticity ^31^. We tested the trends in the cumulative extinctions of species or of interactions during the sequential removal as proportions of the total pollinator richness or of the total interaction quantity with generalized mixed models in the *glmmTMB* package ^32^ in R ^33^. The number of pollinator extinctions (or of interactions) was the response variable, the number of removed plant species was included as a numerical predictor and the observed/TCM/SCM was a categorical one, the total number of pollinators (or of interactions) was an Offset term ^34^; site identity was used as a random intercept.

In addition, we also recorded the amount of species extinctions predicted by the models that were true positives (predicted extinctions which happened in the observed networks), false positives (predicted extinctions which did not happened), true negatives (extinctions not predicted which did not happen in the observed networks) or false negatives (extinctions not predicted which did happen in reality) with both TCM and each SCM simulation at each plant removal stage.

### Networks indices and beta-diversity components

We assembled interaction matrices for each stage of the experiment in all sites and calculated several network-level indices that describe different aspects of species interactions: the binary *Connectance*, indicates the proportion of realised links in relation to all possible links (range 0-1); the weighted *Nestedness NODF* (Nestedness based on Overlap and Decreasing Fill) quantifies the tendency of generalist species to interact with both other generalists and with specialists and ranges 0-100 (the maximum is for fully nested networks); the weighted *Modularity* measures the interactions partitioning into groups, it was computed by the algorithm DIRTLPAwb+ and ranges 0-1 (the maximum is for full compartmentalization); the weighted *H_2_’* measures specialization considering the diversity of interactions based on Shannon entropy and ranges 0-1 (the maximum is for perfect specialisation). The following species-level indices were also calculated: the weighted *Connectivity* and *Participation*, which express the ability of a species to connect partners of different modules (*Connectivity*) or to interact with species of the same module (*Participation*). All these indices were calculated with the *rnetcarto* and *bipartite* packages for R ^35,36^. In addition, an index of network robustness that we name *Stochastic robustness* was calculated as the area under a curve drawn from the rate of pollinators surviving a sequential removal of all plants from the most generalist to the most specialist as simulated by 10^3^ SCM ^10^; this was drawn as a mean number of pollinator extinctions across simulations and was calculated separately for each experimental plant removal phase.

We quantified the turnover of interactions across the removal stages using the approach developed in^37^. This method quantifies the total interaction turnover as *β*_*WN*_ = *β*_*ST*_ + *β*_*OS*_ and partitions it into species turnover (i.e., β_*ST*_, the interaction diversity in the pool of species that are not shared between two networks) and interactions rewiring (i.e. β_*OS*_, switching of interacting partners in species occurring in both networks). These were calculated for all sites and consecutive stages of the experimental removal (before - 1 sp. removed, 1 sp. removed – 2 spp. removed, and so forth) with Whittaker’s beta-diversity index and its components extracted with the package *betalink* ^38^. Values for these indices range from 0 to 1; higher values indicate higher turnover or rewiring.

Two types of interaction matrix were used for the turnover analyses; one, as in^37^, uses binary matrices and focuses on the number of interaction links per species. In addition, to account for the frequency of interactions, we also employed a quantitative version of beta-diversity that is calculated as above but in which the sum of interaction frequency per species is used instead of the number of links.

The effects of plant species removal on network indices and on beta-diversity components were tested with generalized linear mixed models (GLMM) with the *glmmTMB* package in the R environment ^32^; a given index was the response, the site identity was a random intercept, and Beta or Gaussian distributions were used depending on the response variable. For the beta-diversity components, pairs of successive removal stages were used as categorical predictor variables. For the network indices, the number of removed plants was used as numerical predictor. As in ^39^, network size (*AxP*, with *A*=the number of animal species and *P*= the number of plant species in the matrix) and the number of network interactions (the quantitative matrix sum) were included in the models in order to account for their effects on index variation over the experiment. We favoured this approach rather than the delta- or z-transformations because those can cause biases ^40^ and they are more useful for testing departures from a random expectation ^28^, while we aimed at testing the effect of a treatment in causing specific trends (i.e. increase or decrease of an index). To compare the trend of a given index with that of the control, the values from the control site during the experiment were included as an Offset term in the GLMM.

For *Connectivity* and *Participation*, plants and pollinators were analyzed separately in GLMMs with a given index as a response variable, the number of removed plant species as numerical predictor and species identity within site as the random intercept. Here, it was not possible to include the control site for direct comparison because not all species were shared with the removal sites.

### Drivers of interactions

For each site and for each plant removal stage, several simulation models were constructed from different probability matrices to explore the factors driving the observed interactions and indices. The matrices used for the models were: NULL(N) explores the effect of randomness and all species have the same probability of interactions (=1); ABUNDANCES investigates the role of abundances of either or both network levels and the matrix is filled with either the number of flowers of a plant(P), or the abundance of the pollinator species calculated as total abundance of flower visitors of that species over the entire study period (I), or the element-wise multiplication of these two *A=PxI*; (3) in MORPHOLOGICAL-MATCH(M), the matrix is filled with 1 only when a morphological match between length of insect mouthparts and a flower’s nectar allocation depth occurs (Stang *et al.* 2009), for example an insect’s “long-mouthparts” with a flower’s “hidden-nectaries”, “intermediate mouthparts” with “semi-hidden nectaries” and “short-mouthparts” with “accessible nectaries”. As in ^41^, insects were categorized as having a long tongue (>9 mm), intermediate tongue (4-9 mm) or short (<4 mm), and plants for having nectar hidden in flower structures (e.g. larger Fabaceae and flowers with tubular corolla), semi hidden nectaries (more open tubes, smaller Fabaceae) and accessible nectaries (very short tubes or open flowers); (4) SUGAR-RESOURCES(S) assumes that the probability of interaction is proportional to amount of sugar per flower in the nectar and the matrix is filled with the amount of sugar/flowers per plant species ^21^; these values were obtained as detailed in ^25^ by sampling nectar from flowers bagged for 24h and with high performance anion exchange chromatography. For each matrix, probabilities were obtained by dividing the cells of the matrices by the matrix sum. In addition, the interactions of these drivers were included by building models based on multiplying two or three of the matrices described above, specifically: *AxM*, *AxS*, *MxS*, and *AxMxS*. We ran 10^3^ simulated networks with the *mgen* function of the *bipartite* R package that distributes the interaction quantities of the real networks according to the probabilities of the model matrix. For each simulated network, network indices and beta-diversity components were calculated as for the real networks (see above). A given driver is considered as consistent with the empirical observations when its 95% confidence interval includes the real network index ^24^.

To investigate which of the above drivers provided the best fit in terms of predicting the occurrence and frequency of the species pairwise interactions in the observed networks, we used a likelihood approach. Following ^24^, a multinomial distribution was calculated from the interaction frequencies of the observed network and from a given probability matrix. Then, the delta of the Akaike information criteria (ΔAIC) was used to evaluate the ability of each probability model to predict the likelihood of pairwise interactions. As in^42^, in the AIC calculation, the number of parameters was set as the number of species in each probability matrix multiplied by the number of matrices used in order to weight each model’s complexity.

## Results

The plant-flower visitor networks of the experimental sites were similar in species richness (plants *P*=28, pollinators *I*=157 in Site1; *P*=24, *I*=171 in Site2; *P*=20, *I*=106 in Site3).

### Species co-extinctions

The cumulative proportion of observed and predicted extinctions increased linearly with the number of removed plants for both species and interactions (β_species_= 0.158, likelihood ratio test χ^2^_plant removal_ = 176.356, df=1, p<0.001; β_interactions_ = 0.178, likelihood ratio test χ^2^ = 3838.7, df=1, p<0.001, Fig. 1). The observed networks (OBS) registered more species extinctions than the TCM and the SCM models (β_OBS-TCM_=1.03, β_OBS-SCM_=0.958, likelihood ratio test χ^2^_OBS/TCM/SCM_ =42.493, df=1, p<0.001). Similarly, the observed networks lost more interactions than what was predicted by the two models (β_OBS-TCM_= 0.9, β_OBS-SCM_= 0.713, likelihood ratio test χ^2^_observed/TCM/SCM_ = 359.1, df=1, p<0.001).

**Figure 1.**
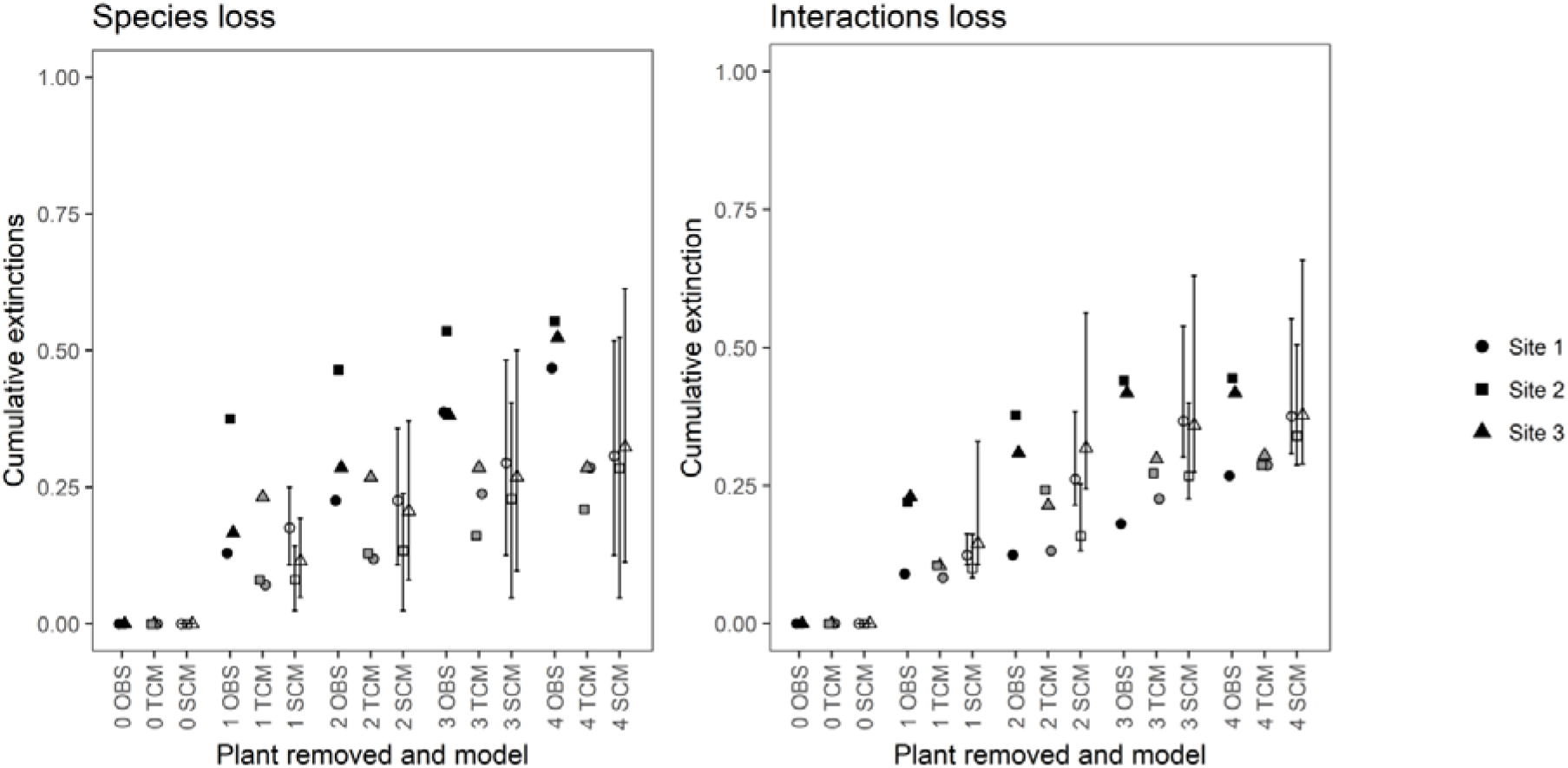
Species coextinctions. Cumulative proportion of extinctions of species and of interactions over the sequential plant removal as observed in the real networks (“OBS”) and as predicted by TCM and SCM co-extinction models for each site. In SCM, the symbols and lines indicate the mean and 5%-95% quantiles of 10^3^ simulations. Statistical tests are presented in the Results.

Among pollinator species which went extinct in the field experiment, on average 85.33% (range across sites and plant removal stages: 33 – 100%) were species which were predicted to go extinct by the TCM (true positives), while the rest were species which the model incorrectly predicted to survive (false positives). SCM provided worse predictions of extinctions of individual species with the mean of 26.62 % true positives (range: 10 – 37.7 %). On the other hand, among species which survived in the field experiment, on average 33.66% were species predicted to survive based on TCM (range 15 – 50%) and 41.29 % based on SCM (range 10.88 – 85.53%) (true negatives), the rest were extinctions observed in the field, but not predicted by the models (false negatives).

### Network structure and interaction turnover during plant removal

Network modularity and specialization significantly increased with the number of removed plants, while nestedness decreased significantly (Fig. 2, Table 1). However, when the values from the control site were used as an offset, the statistical significance of the increase in modularity and decrease in nestedness was confirmed, but the significance of specialization was not confirmed (Table 1). The trends of other network indices were not significant during the sequential plant removal. In the species-level indices, plants and pollinators responded differently (Fig. 2, Table 1). Only the plant Connectivity increased significantly, while plant Participation and the pollinator indices were nearly constant during the sequential plant removal.

**Table 1.** Statistics of the network indices and interaction turnover components. Each row is a separate generalized mixed effect model, further details are described in Methods. WN is total beta diversity, OS is the rewiring of interactions, ST is the species turnover between networks pairs. “W.” stands for “weighted”. Δ*AIC* is calculated as *AIC_i_ - AIC_min_*. Statistically significant predictors (P<0.05) are highlighted in bold. Significance of the models including the values of the indices in the control site as an offset are also given.

| | Df | $\Delta AIC$ | $\chi^2$ | P | P with control offset |
| --- | --- | --- | --- | --- | --- |
| Connectance | 1 | 1.274 | 0.726 | 0.394 | 0.719 |
| NODF | 1 | 6.032 | 8.032 | **0.005** | **0.001** |
| Modularity | 1 | 7.246 | 9.246 | **0.002** | **0.007** |
| H2' | 1 | 11.076 | 13.076 | **<0.001** | 0.073 |
| Stochastic robustness | 1 | 1.819 | 3.819 | 0.051 | 0.350 |
| Connectivity plants | 1 | 10.439 | 12.439 | **<0.001** | NA |
| Participation plants | 1 | 1.857 | 0.143 | 0.705 | NA |
| Connectivity pollinators | 1 | 5.285 | 7.285 | **0.007** | NA |
| Participation pollinators | 1 | 1.509 | 0.491 | 0.484 | NA |
| WN ($\beta$) | 3 | 0.604 | 5.396 | 0.145 | 0.110 |
| OS (rewiring) | 3 | 3.793 | 2.207 | 0.531 | 0.525 |
| ST (turnover) | 3 | 2.262 | 3.738 | 0.291 | 0.323 |
| W. WN ($\beta$) | 3 | 4.170 | 1.830 | 0.608 | 0.688 |
| W. OS (rewiring) | 3 | 2.581 | 3.419 | 0.331 | 0.698 |
| W. ST(turnover) | 3 | 1.400 | 4.600 | 0.204 | 0.890 |

**Figure 2.**
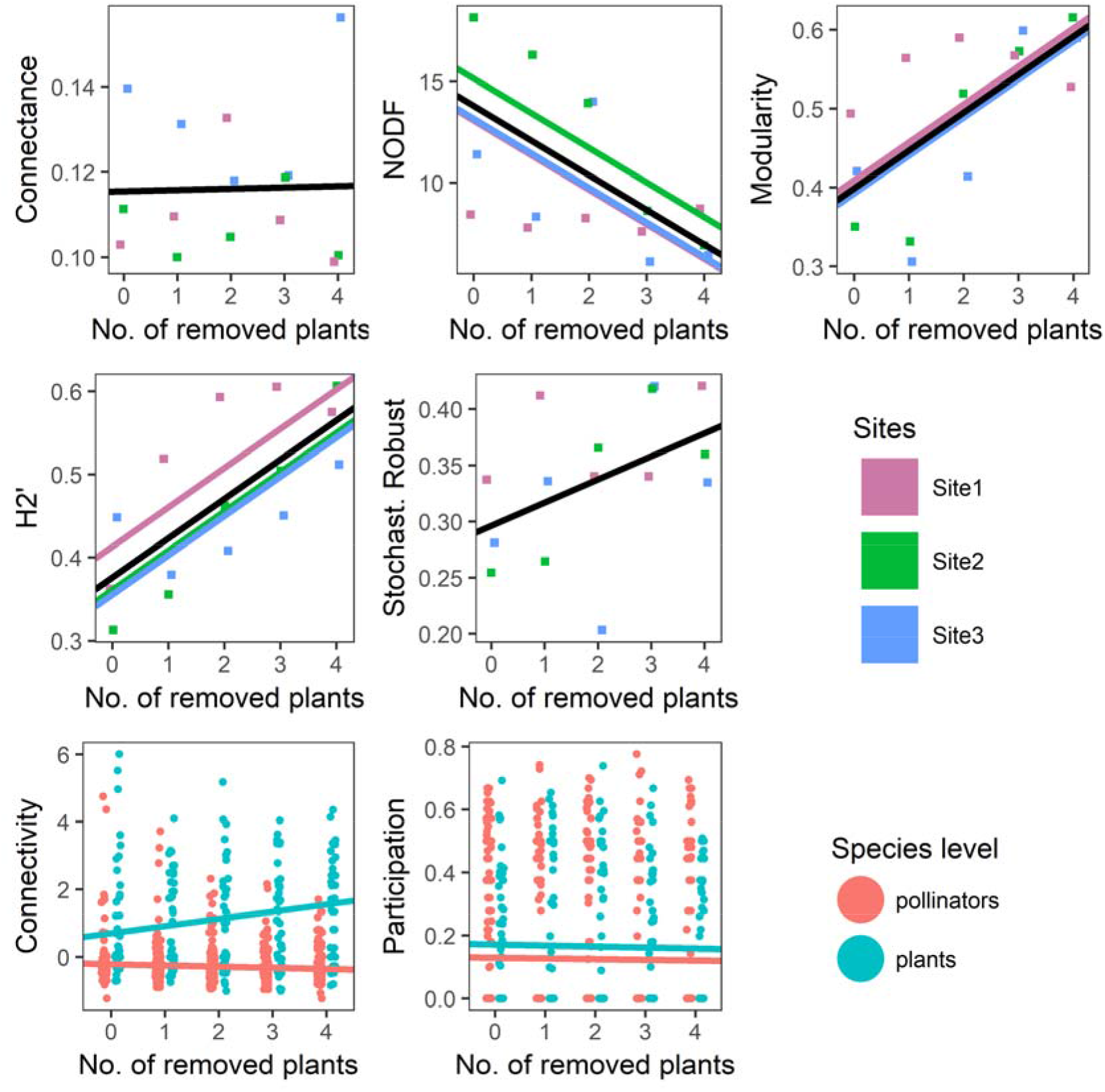
Network structural changes. Network indices responses to removal of generalist plants. The significances of predictors are expressed in Table 1. The black line is the average trend predicted by the models, coloured lines are predictions for each site; plots with black line only are cases which were not statistically significant.

The interaction turnover was high in both quantitative and binary versions (Fig. 3), with a larger proportion attributable to rewiring than to species turnover; however, no statistically significant trend was found in these indices in response to the treatment (Table 1).

**Figure 3.**
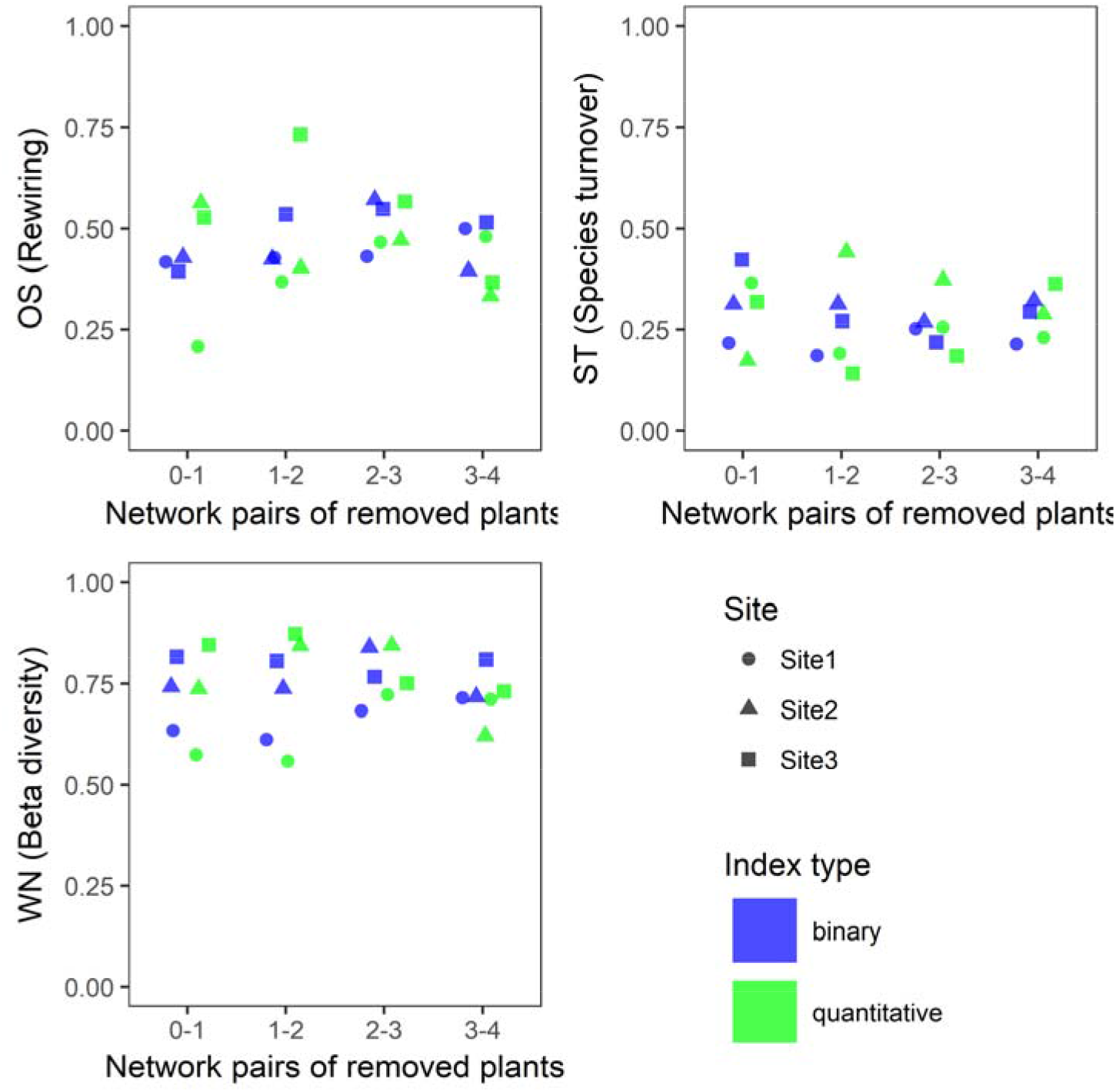
Species interaction rewiring. Interaction turnover of pairs of networks after each stage of plant removal. Both the binary indices and their quantitative counterparts are plotted. Significances of predictors are included in Table 1.

### Drivers of network structure and interaction turnover

In the likelihood analysis (Table 2), models based on species abundance usually provided the best fit to the observed species interactions, especially in the case of the pollinator abundances model (*I*); the null model assuming equal probability of all interactions (NU) predicted the observed interactions as the plant removal progressed; the *S* model based on the amount of sugars in nectar (*S*) also contributed to describing the interactions (i.e. it had low ΔAIC values).

**Table 2.**
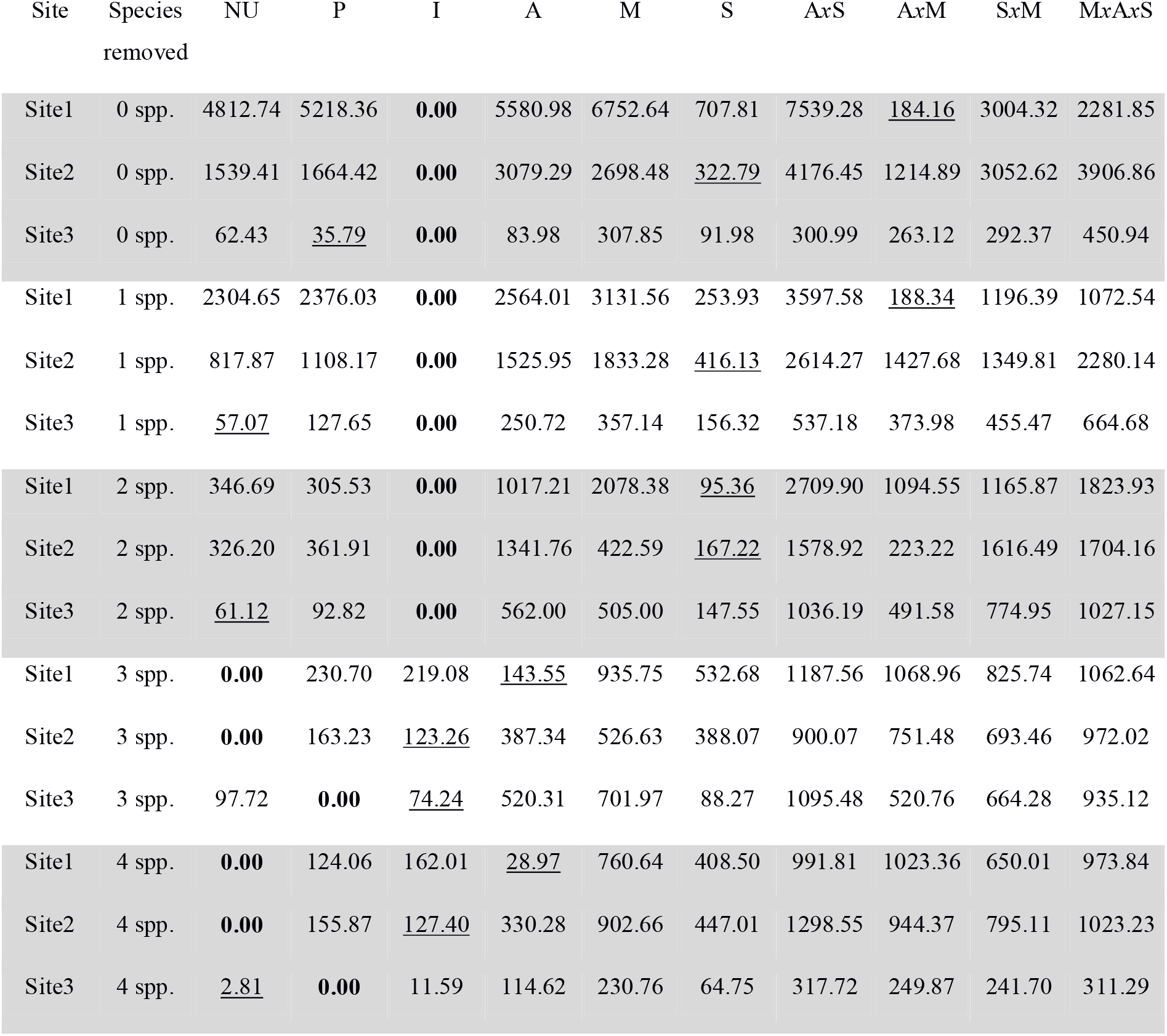
Models’ likelihood of pairwise species interactions drivers (Δ*AIC*). In bold, the probability models that best predicted the interactions are highlighted; the second important probability models are underlined. Model acronyms are described in the Methods.

| Site | Species removed | NU | P | I | A | M | S | AxS | AxM | SxM | MxAxS |
| --- | --- | --- | --- | --- | --- | --- | --- | --- | --- | --- | --- |
| Site1 | 0 spp. | 4812.74 | 5218.36 | **0.00** | 5580.98 | 6752.64 | 707.81 | 7539.28 | <u>184.16</u> | 3004.32 | 2281.85 |
| Site2 | 0 spp. | 1539.41 | 1664.42 | **0.00** | 3079.29 | 2698.48 | <u>322.79</u> | 4176.45 | 1214.89 | 3052.62 | 3906.86 |
| Site3 | 0 spp. | 62.43 | <u>35.79</u> | **0.00** | 83.98 | 307.85 | 91.98 | 300.99 | 263.12 | 292.37 | 450.94 |
| Site1 | 1 spp. | 2304.65 | 2376.03 | **0.00** | 2564.01 | 3131.56 | 253.93 | 3597.58 | <u>188.34</u> | 1196.39 | 1072.54 |
| Site2 | 1 spp. | 817.87 | 1108.17 | **0.00** | 1525.95 | 1833.28 | <u>416.13</u> | 2614.27 | 1427.68 | 1349.81 | 2280.14 |
| Site3 | 1 spp. | <u>57.07</u> | 127.65 | **0.00** | 250.72 | 357.14 | 156.32 | 537.18 | 373.98 | 455.47 | 664.68 |
| Site1 | 2 spp. | 346.69 | 305.53 | **0.00** | 1017.21 | 2078.38 | <u>95.36</u> | 2709.90 | 1094.55 | 1165.87 | 1823.93 |
| Site2 | 2 spp. | 326.20 | 361.91 | **0.00** | 1341.76 | 422.59 | <u>167.22</u> | 1578.92 | 223.22 | 1616.49 | 1704.16 |
| Site3 | 2 spp. | <u>61.12</u> | 92.82 | **0.00** | 562.00 | 505.00 | 147.55 | 1036.19 | 491.58 | 774.95 | 1027.15 |
| Site1 | 3 spp. | **0.00** | 230.70 | 219.08 | <u>143.55</u> | 935.75 | 532.68 | 1187.56 | 1068.96 | 825.74 | 1062.64 |
| Site2 | 3 spp. | **0.00** | 163.23 | <u>123.26</u> | 387.34 | 526.63 | 388.07 | 900.07 | 751.48 | 693.46 | 972.02 |
| Site3 | 3 spp. | 97.72 | **0.00** | <u>74.24</u> | 520.31 | 701.97 | 88.27 | 1095.48 | 520.76 | 664.28 | 935.12 |
| Site1 | 4 spp. | **0.00** | 124.06 | 162.01 | <u>28.97</u> | 760.64 | 408.50 | 991.81 | 1023.36 | 650.01 | 973.84 |
| Site2 | 4 spp. | **0.00** | 155.87 | <u>127.40</u> | 330.28 | 902.66 | 447.01 | 1298.55 | 944.37 | 795.11 | 1023.23 |
| Site3 | 4 spp. | <u>2.81</u> | **0.00** | 11.59 | 114.62 | 230.76 | 64.75 | 317.72 | 249.87 | 241.70 | 311.29 |

In the networks and beta-diversities (Fig. 4, and Supplementary Fig. S1, S2, S3, S4), none of the models generated CI including every observed index. The *Connectance* and *H2’* were particularly poorly predicted. Remarkably, both *P* and *I* models and the multiplication of abundances with others were explaining the observed indices in multiple cases (e.g. *NODF, Modularity, WN, OS, ST*) and *MxS* predicted *NODF* in most cases. In addition, the predictors usually changed as the removal of plants progressed, such as *M* and *S* predicting both *OS* and *ST* after the first removal, and, before removal, both *M* predicting the *Modularity* and *M* with abundances predicting *H2’*. In some cases, the complexity of the models (i.e. from the multiplication of probability matrices) either increased the predicting power (*Modularity*, *NODF, WN, OS*) or decreased it (*H2’*, *ST*) as the removal progressed.

**Figure 4.**
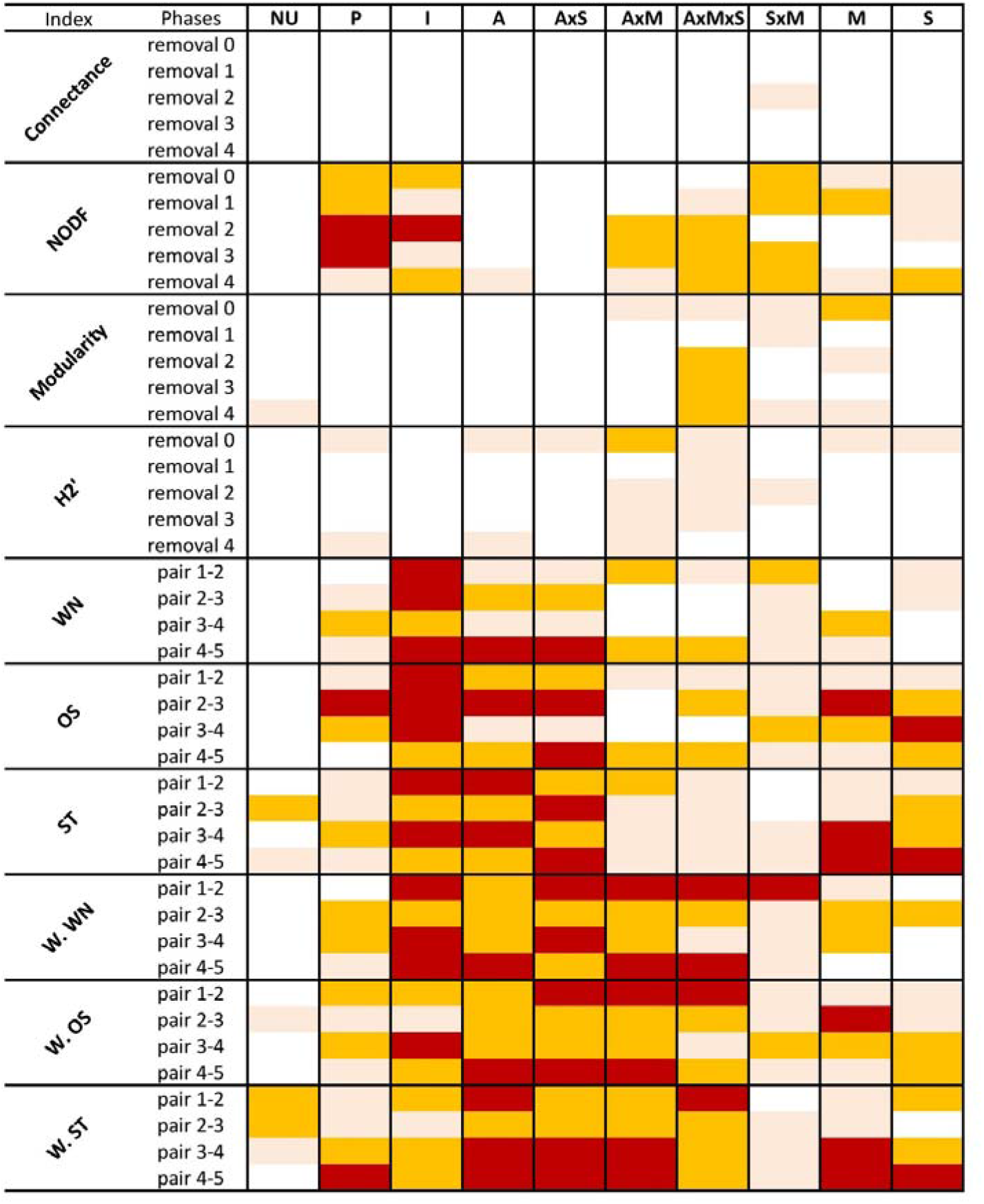
Ecological drivers of network indices and rewiring. Heatmap indicating the overlap between the observed network or beta-diversity indices and the confidence intervals of 10^3^ simulations generated from probability matrices (in columns; model acronyms are described in the Methods.). Colours symbolize the number of sites being predicted correctly: red is for correct prediction in 3 sites, orange is for 2 sites and pink is for 1 site.

## Discussion

In our study, after removing generalist plants from real plant pollinator networks, both species and interaction extinctions increased more than expected from co-extinction models. As far as we know, previous studies have only used *in-silico* estimation of extinctions ^8,33,34^, but our experiment clarifies that TCM and SCM underestimated species extinction rates, and the rate of false positives and false negatives was high in relation to the identity of the species that were lost. Furthermore, these models underestimated the rate of interaction loss, which is a major flaw of currently available simulations, an issue that has been already pointed out ^11^. Altogether, field experiments such as the one we performed have a big potential for validating, rejecting or refining the theoretical insights gained by simulation models, and could trigger further development of more accurate models on network functioning, stability and co-extinction rates. We speculate that the differences between predicted and observed extinctions of this study could be due to ecological factors not accounted for by the coextinction models^35^. The two simplest possible explanations might be that, firstly, the treated sites became progressively less attractive to foraging pollinators which emigrated even when resources they were using were still available, because pollinators are attracted by total flowering plant richness and abundance (e.g. ^47^). Another possible explanation is that plant removal made the network structure became more fragile, so that species would became less anchored to other species in the interaction web and more exposed to extinction ^37,38^.

In our study, the removal of generalist plants clearly impacted network structure. The structure of a plant-pollinator network after a disturbance can provide information on its functionality thanks to the ecological interpretation of network indices ^30,31^. For example, networks usually organize in nested patterns where generalists interact with both specialists and other generalists, but the decrease of nestedness after a perturbation could indicate a loss of interactions mainly affecting the specialist species^32^. In our study, the loss of generalist plants triggered a decrease of nestedness and an increase of modularity. These trends could relate to the fact that specialisation also increased during the successive plant species removal, possibly as a result of decreased pollinator abundances as previously shown by ^26^, i.e. reductions in the number of interactions triggers changes in network structure ^39,40^. Specifically, the observed decrease of nestedness is a possible symptom of instability ^5^ because specialist species are less connected to the generalist network core ^30^. Concurrently, the observed increase in modularity indicates an emergence of a compartmentalised structure, a sign of potential network breakdown ^3^. Although compartmentalization of predator-prey food-webs is considered beneficial as it buffers against alterations spreading throughout the entire web ^41^, in mutualistic networks a very high modularity actually prevents the access to alternative resources and it can be linked to decreased stability ^28^. Therefore, these results seems to support the expectation that after removing key elements, the network shifts towards a less cohesive structure fragmented in compartments. Changes in nestedness and modularity did not translate into a lower stochastic robustness index. A possible reason for this relies on the dynamic yet asymmetric re-organization of species interactions along the sequence of plant removal. While the remaining plant species became increasingly centralized in the network, there was no trend in the average centralization of pollinator species. In addition, the network rewiring was high and played a larger role than species turnover in determining interaction turnover during the experiment, as in ^16^, although without a clear trend during the experiment. As rewiring has traditionally been associated with network stability ^19,31^, both the observed increased plant centrality and high rewiring could explain why network robustness index increased as more plants were removed.

Many indices of the studied networks were explained by complex combinations of predictors, such as the interaction of abundances with morphological match and with sugar rewards. This was especially the case as several plants were removed, which could suggest a prominence of network complexity following the removal of generalist plants. Conversely, simpler models (i.e., based on only one predictor) explained only occasionally some of the indices. Some remarkable examples are: flower abundance predicting nestedness, which reflects the role of abundant generalist plants which interact with numerous insects and drives the nested pattern ^42^; modularity being predicted, before plant removal, by the morphological match of corolla depth and mouthpart length, suggesting that trait matching is relevant in defining modules ^21,43,44^; nestedness being predicted by the interaction of morphological matching and sugar amount in the nectar, which confirms that trait matching allows an efficient resource gathering ^45,46^; network rewiring being predicted by abundances, and by the interaction of abundances and sugar amount, as the abundant and rewarding plants are more central in the network and more likely to establish new interactions ^47,48^; morphology matching predicting the rewiring in some stages of plant removal, indicating that utilization of new resources is constrained within trait-spaces ^26^.

On the other hand, individual pairwise interactions were explained best by the model using pollinator abundances, during the entire experiment. However, when several plants were removed, pairwise interaction were instead explained by the null model assuming equal probability of interactions. While the role of pollinator abundance reflects the relationship between abundance and generalization of interactions ^24,25^; the superior fit of the null model in the later stages of the experiment suggests an emergence of randomness in species interactions in impoverished communities. The role of randomness in ruling pairwise interactions is particularly alarming, because it would indicate the disruption of established interaction assembly mechanisms, and may also be linked to opportunism in interactions and high rewiring ^27^.

Since the experiment stopped when four species were removed, we do not know if the observed linear trends in network indices and species and interaction extinctions would also progress linearly when the other remaining plants are taken away from the system. Theoretical studies have used various approaches, with models assuming linearity ^49^ and non linear models ^50^, which could generate conflicting results ^51^. Nevertheless, some theoretical studies consider linear functional responses as uncommon trends in mutualistic interactions (e.g. ^66^). Thus, it could be expected that, when other additional plant species are removed, the trend would become nonlinear, for instance as observed in co-extinction models ^6^.

In conclusion, pollinators occurrence and species interactions are more sensitive to the disappearance of generalist plants than is expected from network co-extinction models. When the key plants are removed, the network structure changes, extinctions of species and interactions increase, and opportunism can become more prominent. This gives strong support to proposals indicating that conservation of interactions should be centred on the generalist species ^29,53^. However, this generalist-based conservation view should consider the dynamics and re-organization of interactions and the asymmetrical responses between plants and pollinators, which compensate for an even more detrimental collapse of species networks.

## Acknowledgements

The authors thank Vojtech Novotny and Darren Evans for their valuable comments on this study. The authors would like to thank Jana Jersáková, Štěpán Janeček, Dagmar Hucková, Tomáš Gregor, Michal Rindoš, Michal Bartoš, Zuzana Chlumská for their help during laboratory and field work. We thank also the specialists who identified some of the insects collected: Daniel Benda, Jiří Beneš, Jiří Hadrava, Petr Heřman, Irena Klečková, Petr Kment, Oldřich Nedvěd, Hana Šuláková, Michal Tkoč, and Šimon Zeman. This project was supported by the Czech Science Foundation (projects GP14-10035P and GJ17-24795Y) and AA was also supported by a grant GA JU 152/2016/P provided by the University of South Bohemia. The funders had no role in study design, data collection and analysis, decision to publish, or preparation of the manuscript.

## Author Contribution

Data collection: PB, JK, AA. Funding acquisition: JK. Methodology: PB, AA, JK, JO, AN. Statistical analyses: PB. Supervision: JK, JO, AN. Visualization: PB. Writing original draft & editing: PB, JO, AA, JK.

## Code availability

The script code used for the analyses of this manuscript is available upon request from the corresponding author.

## Data availability statement

All relevant data are within the paper or stored in a public repository (this will be associated to a DOI upon manuscript acceptance)

## Competing interests

The authors have declared that no competing interests exist.

## Ethics statement

No permits were required for this project.

## Supplementary information

Figure S1. Network Connectance and Nestedness indices predictions by probability models for each site.

Figure S2. Network Specialization and Modularity indices predictions by probability models for each site.

Figure S3. Qualitative interaction turnover indices predictions by probability models for each site.

Figure S4. Quantitative interaction turnover indices predictions by probability models for each site.

