## Supplementary Fig. S1, S2, S3, S4 for "An empirical attack tolerance test alters the structure and species richness of plant-pollinator networks"

### Supplementary information

Figure S1. Network Connectance and Nestedness (W NODF) indexes detection by probability models (NU= null, P= plant abund., I= Insect abund., A= abund., M= morphological match, S= sugar amount). The vertical lines represent the observed values (solid line for Site 1, dashed for Site 2, loosely dashed for Site3). The horizontal segments are the ranges from the simulations and are vertically distributed (Site 1, under Site 2 that are under Site 3).

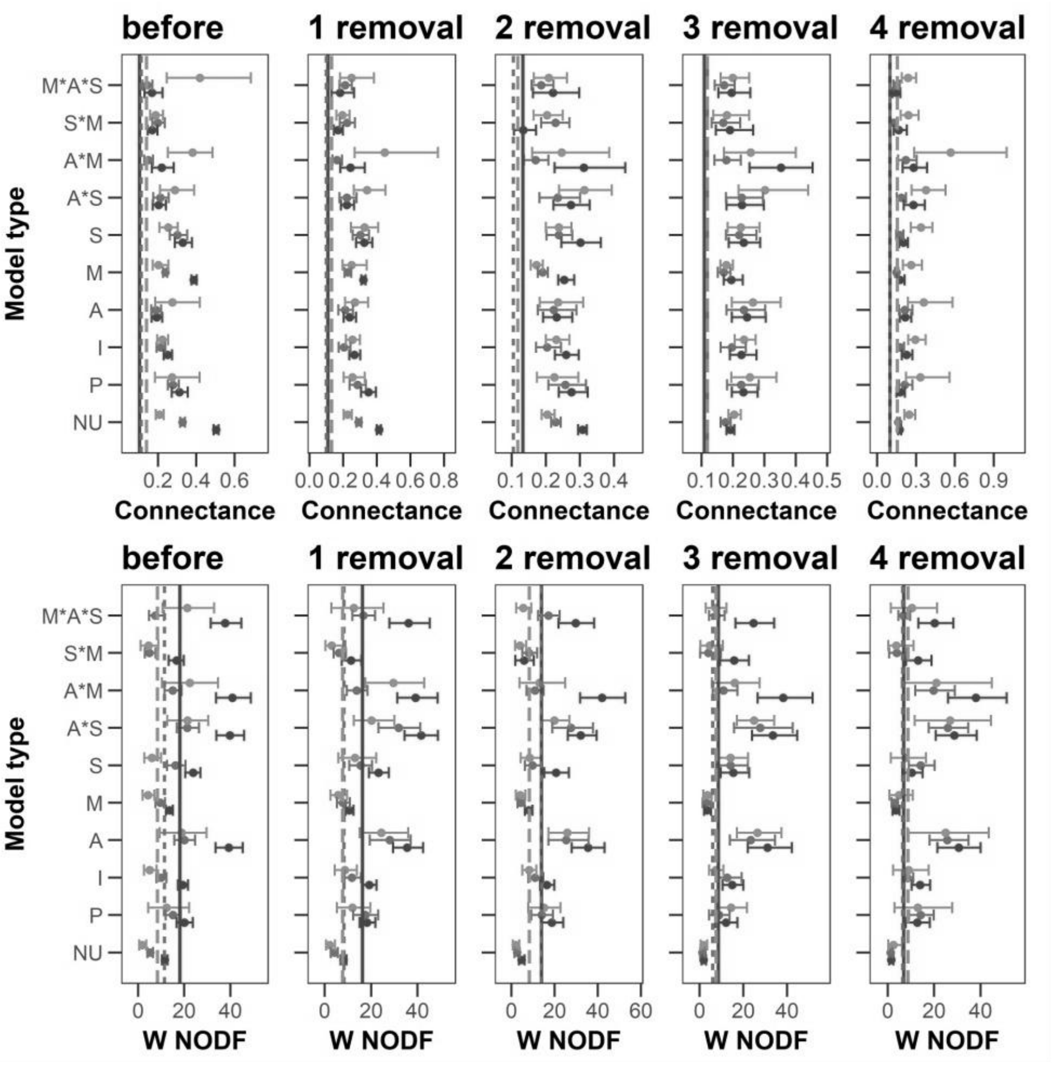

Figure S2. Network Specialization (W. H2') and Modularity indexes detection by probability models (NU= null, P= plant abund., I= Insect abund., A= abundances, M= morphological match, S= sugar amount). The vertical lines represent the observed values (solid line for Site 1, dashed for Site 2, loosely dashed for Site3). The horizontal segments are the ranges from the simulations and are vertically distributed (Site 1, under Site 2 that are under Site 3).

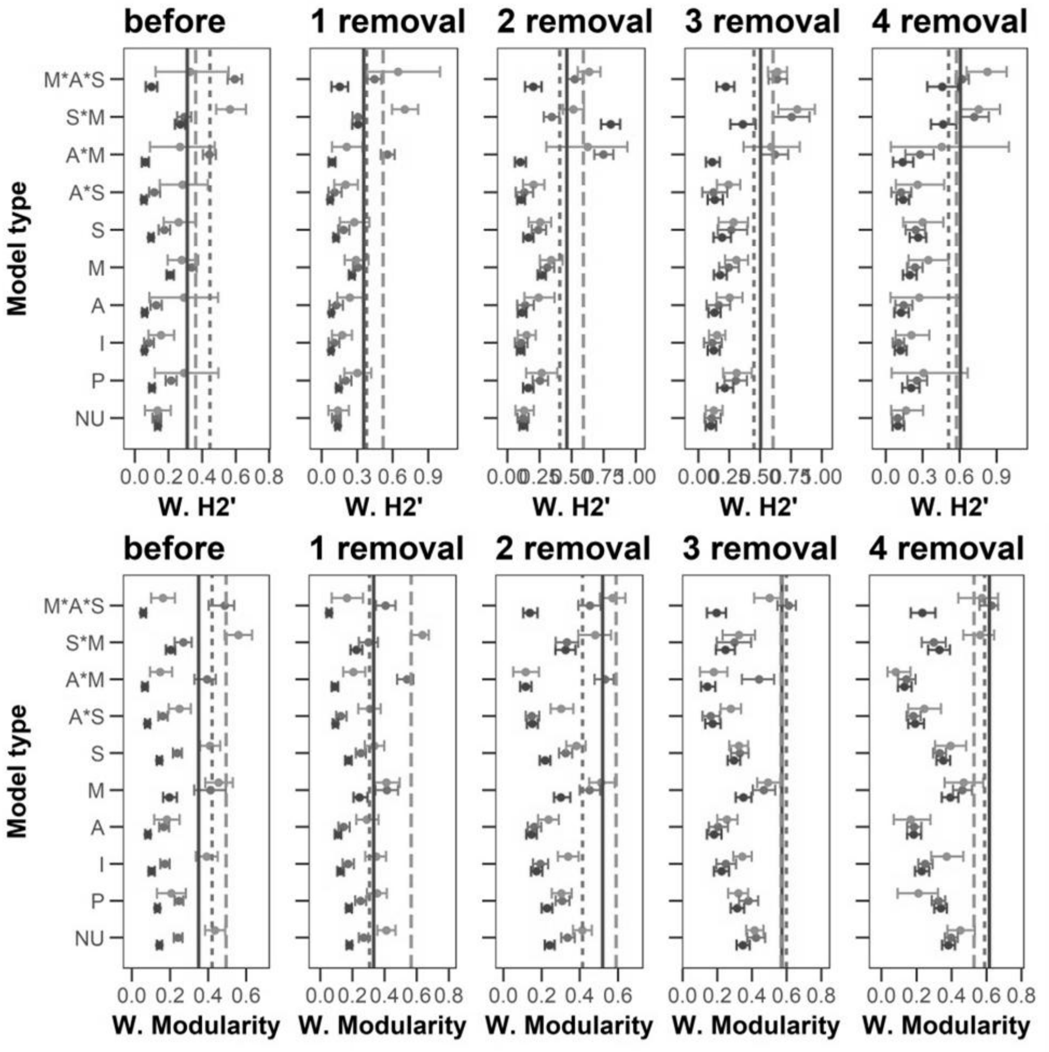

Figure S3. Qualitative interaction turnover indexes detection by probability models (NU= null, P= plant abund., I= Insect abund., A= abund., M= morphological match, S= sugar amount). The vertical lines represent the observed values (solid line for Site 1, dashed for Site 2, loosely dashed for Site3). The horizontal segments are the ranges from the simulations and are vertically distributed (Site 1, under Site 2 that are under Site 3).

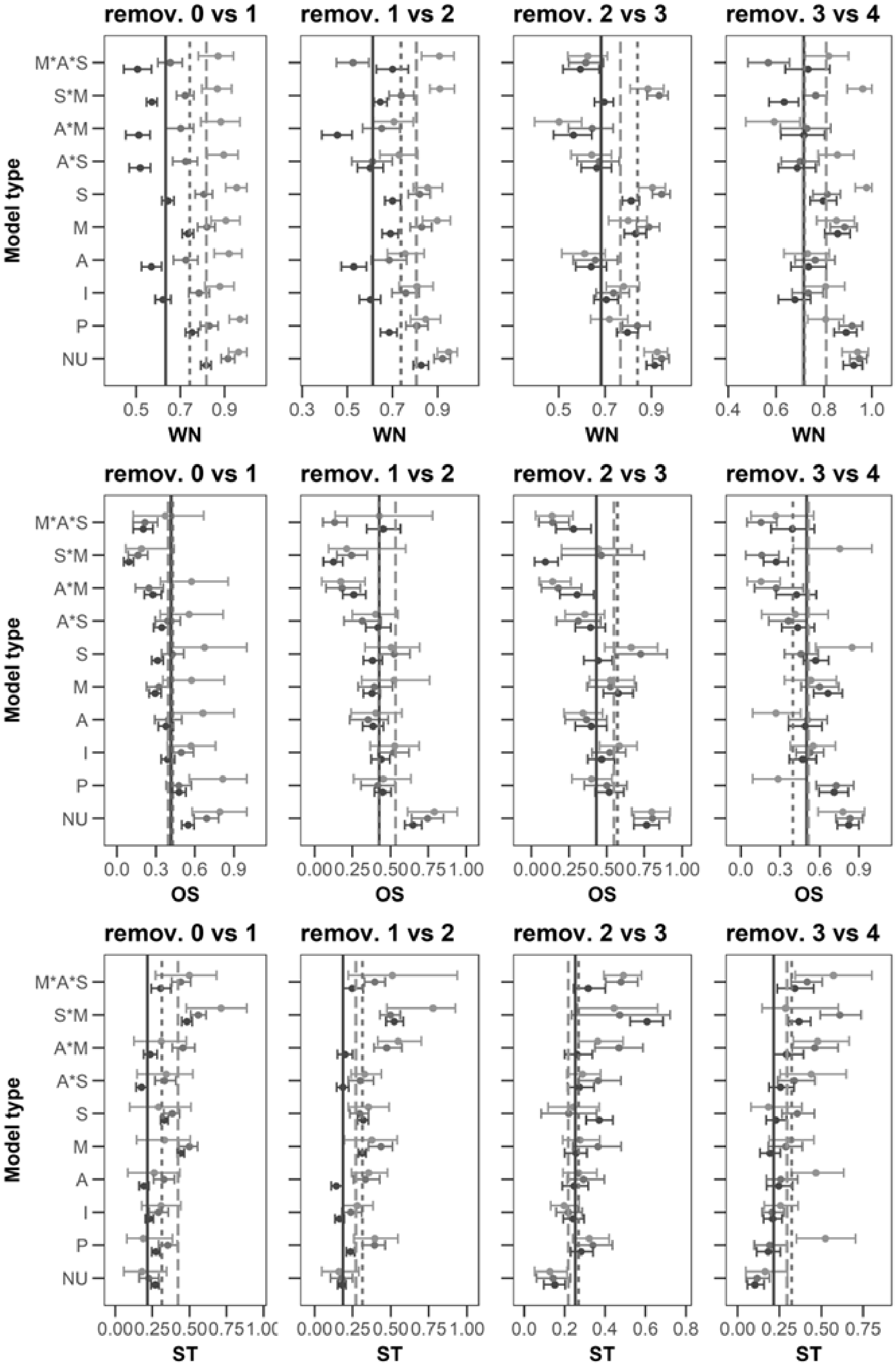

Figure S4. Quantitative interaction turnover indexes detection by probability models (NU= null, P= plant abund., I= Insect abund., A= abund., M= morphological match, S= sugar amount). The vertical lines represent the observed values (solid line for Site 1, dashed for Site 2, loosely dashed for Site3). The horizontal segments are the ranges from the simulations and are vertically distributed (Site 1, under Site 2 that are under Site 3).

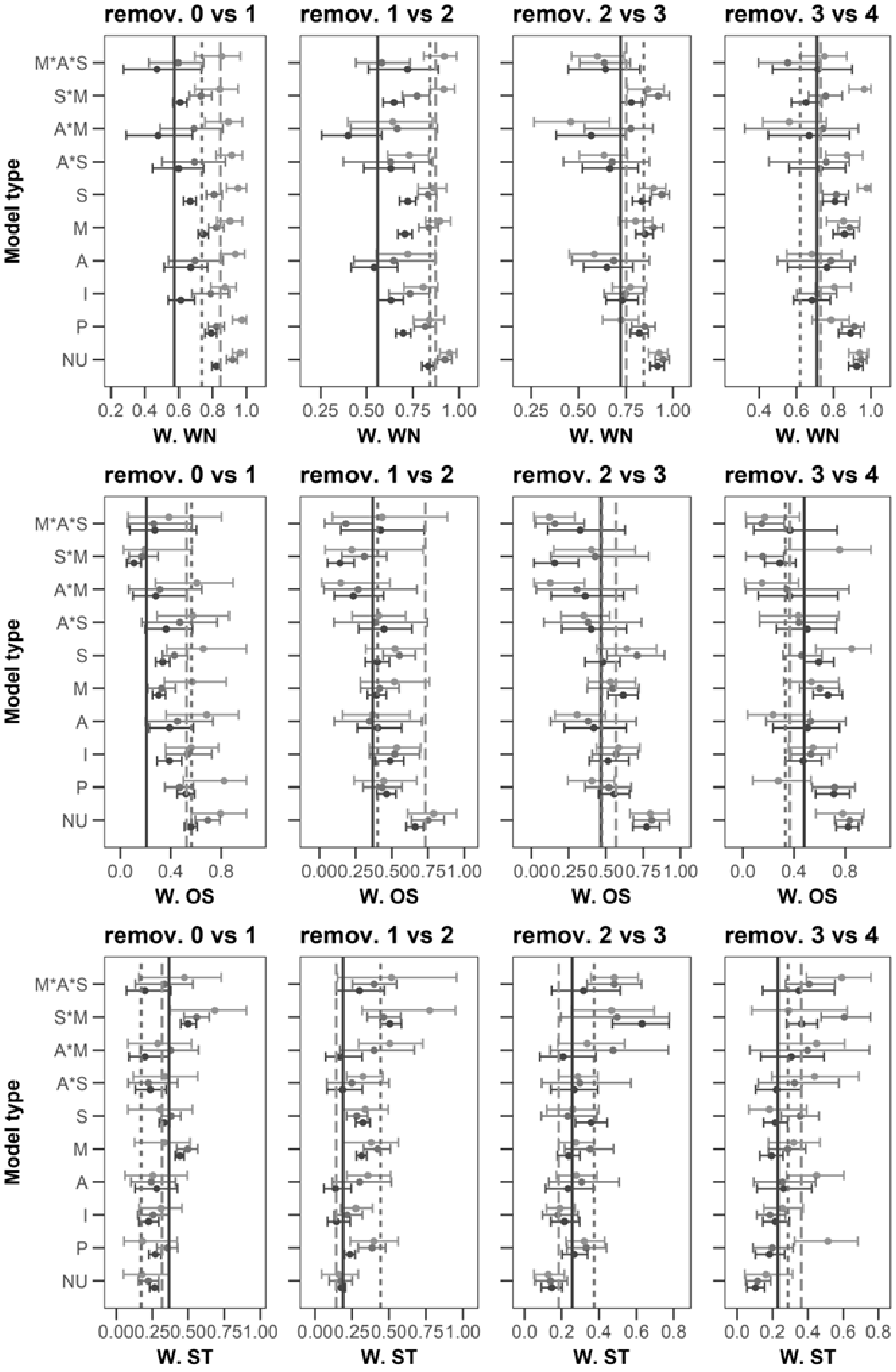
